# Multidimensional brain-age prediction reveals altered brain developmental trajectory in psychiatric disorders

**DOI:** 10.1101/2020.12.24.424350

**Authors:** Xin Niu, Alexei Taylor, Russell T. Shinohara, John Kounios, Fengqing Zhang

**Affiliations:** Department of Psychology, Drexel University, 3201 Chestnut Street Philadelphia, PA 19104, USA; Center for Biomedical Image Computation and Analytics, University of Pennsylvania, Philadelphia, PA 19104, USA; Penn Statistics in Imaging and Visualization Center, Department of Biostatistics, Epidemiology and Informatics, University of Pennsylvania, Philadelphia, PA 19104, USA

**Keywords:** brain-age prediction, multimodal neuroimaging, machine learning, brain development, psychiatric disorders

## Abstract

Neuroimaging-based brain-age prediction has emerged as an important new approach for studying brain development. However, brain regions change in different ways and at different rates. Unitary brain-age indices used in previous studies represent developmental status averaged across the whole brain and therefore do not capture the divergent developmental trajectories of various brain structures. Importantly, this staggered developmental unfolding, determined by genetics and postnatal experience, is implicated in the progression of psychiatric and neurological disorders. Here we propose an analytic method for computing a multidimensional brain-age index (MBAI) that provides regional age predictions. Using a database of 556 subjects (ages 8-21) that includes psychiatric and neurological patients as well as healthy controls, we conducted robust regression and cluster analyses to identify clusters of imaging features with distinct developmental trajectories. We then built machine-learning models to obtain brain-age predictions from each of the identified clusters to form the MBAI. Our results show that the MBAI provides a flexible analysis of region-specific brain-age changes that are invisible to unidimensional brain-age prediction methods. Importantly, brain ages computed from region-specific feature clusters contain complementary information and demonstrate differential ability to classify disorder groups (e.g., specific phobia, depression, ADHD) from healthy controls. Compared to unidimensional brain-age indices, we show that the MBAI is sensitive to alterations in brain structures and captures distinct regional change patterns which may serve as biomarkers that may contribute to our understanding of healthy and pathological brain development and to the characterization, diagnosis, and, potentially, treatment of various disorders.

## Introduction

The human brain shows striking functional and structural changes across the lifespan. On the neuronal level, these changes include synaptic pruning and axonal myelination (1, 2) related to the maturation of cognitive, emotional, and social abilities. On the macroscopic level, brain changes reflect the reduction of gray matter volume (3), cortical thinning (4), and increased integration of structural and functional connectivity (5, 6). Because abnormal changes are associated with symptoms and disrupted cognition in psychiatric disorders, studying them can contribute toward the development of diagnostic and prognostic biomarkers for psychopathology (7) which have the potential to advance our understanding of comorbidity across, and heterogeneity within, mental disorders (8, 9).

Brain regions and systems do not develop at the same rate or in the same way. For example, the reduction of gray matter occurs early in the sensory-motor cortex followed by regions involved in higher-level functions such as the prefrontal and temporal cortex (3). A longitudinal study found a nonlinear brain development pattern of gray matter in the frontal and temporal lobes with peak values emerging at different ages whereas occipital white and gray matter show linear change (10). In addition, many psychiatric disorders emerge during adolescence when dramatic brain changes occur (11, 12). The staggered development of brain regions is both genetically determined and influenced by postnatal experiences (13, 14). Although altered brain development is understood to be associated with psychiatric and neurological disorders such as schizophrenia, mild cognitive impairment, Alzheimer’s disease, and autism (4, 15–17), it is unclear how staggered development is implicated in the progression of psychiatric disorders.

The recent development of brain-based age prediction has emerged as an important approach for quantifying human brain development (18, 19). Machine-learning brain-age prediction based on neuroimaging features has achieved substantial accuracy (20–23). The brain-age gap (BAG), the difference between the chronological age and predicted age based on brain features extracted from neuroimaging data is not interpreted only as noise in the model fitting but as including systematic deviation due to the delay or precocity of brain development. This interpretation is substantiated by the finding that the BAG is associated with a wide range of cognitive and emotional abilities (22, 24).

Despite these promising findings, brain-age prediction still faces challenges. First, the interpretation of the BAG has been inconsistent across studies. Some studies have interpreted it as maturity (22, 24), whereas others have interpreted it as age-related decline (17, 25, 26), even with young participants. Second, machine-learning models can overfit the data and yield nearly perfect prediction accuracy of chronological age which reduces the utility of the BAG as a biomarker for psychiatric disorders (27). As true brain-age is unknown, it is unclear how to reduce noise in the prediction of age while preserving an accurate estimate of the BAG. Third, as different brain systems mature with unique temporal patterns across the lifespan (3, 11, 12), a unidimensional brain-age cannot capture the divergent developmental trajectories of the various brain systems. Specifically, if one part of an individual’s brain is older while another part is younger than average, then the overall predicted brain-age may not show any abnormalities.

To deal with these challenges to brain-age prediction, we developed an analytic pipeline to obtain a multidimensional brain-age index (MBAI) (Fig. 1). We first built a robust regression model (i.e., Huber regression) to predict each brain-imaging feature using chronological age, sex, and their interaction. The estimated regression coefficients characterize the developmental trajectory of each brain-imaging feature. We then conducted cluster analyses on the extracted regression coefficients to identify subgroups of brain-imaging features with a similar developmental trajectory. For each identified cluster, we predicted brain-age via machine learning models (i.e., ridge regression) using imaging features belonging to that cluster. The predictions of brain-age from multiple clusters form the MBAI.

**Figure 1.**
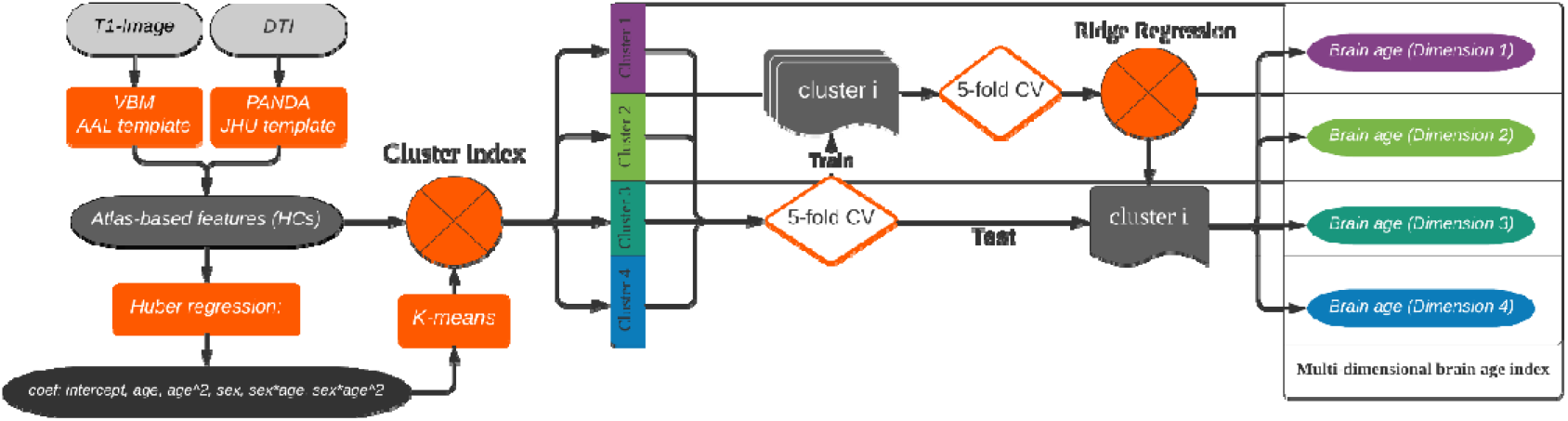
The pipeline of brain imaging processing and multidimensional age prediction. Brain-imaging data from 556 subjects were preprocessed with VBM and the PANDA toolbox for T1 and DTI images, respectively. Atlas-based features were obtained with the AAL template and JHU white-matter template for gray-matter volume (GMV) and fractional anisotropy (FA), respectively. For the HCs, a robust regression model was adopted to model each brain-imaging feature with a quadratic regression model. The coefficients obtained from the regression were fed into a K-means cluster analysis to identify four clusters of brain-imaging features with divergent developmental patterns. For features in each cluster, nested 5-fold cross-validation (CV) was conducted with ridge regression to estimate the brain age for the HCs. The trained model was then applied to the disorder groups to obtain their brain age estimates.

We hypothesize that our proposed MBAI has several advantages compared to the unidimensional one. First, it allows for a more flexible interpretation of the BAG across brain subsystems because it can reveal accelerated development in one subsystem while showing delayed development in another. In contrast, a unidimensional brain-age index captures only the average developmental status across the entire brain. Second, prediction of brain age using a subset of homogeneous features circumvents the curse of dimensionality in brain-imaging data and helps to mitigate overfitting of the age prediction model thereby improving estimates of the BAG. Finally, the MBAI captures distinct growth patterns of the brain systems and is more sensitive to alterations in constrained brain structures. To further validate the clinical utility of our proposed multidimensional brain age index, we showed that the MBAI is altered in several psychiatric disorders. These results demonstrate the utility of the MBAI and support its use as a tool for investigating healthy and disordered brain development.

## Materials and methods

### Datasets

Subjects were selected from the Philadelphia Neurodevelopmental Cohort (PNC) study (28). The PNC research initiative includes a large-scale sample with computerized clinical assessment, neurocognitive testing, and multi-modal brain imaging data. It aims to investigate how brain maturation mediates the development of cognition in healthy people and patients with psychiatric illness (29). Details of subject recruitment and study procedures can be found in published papers (28, 30). All subjects included in this study have associated multimodal brain-imaging data (i.e., T1 weighted MRI and DTI). They also completed a computerized neurocognitive battery (CNB) test (31). After excluding 17 subjects with severe general medical problems, we selected 166 healthy controls (HCs) and 6 groups of participants with psychiatric disorders including specific phobia (N=70), social phobia (N=77), PTSD (N=44), depression (N=70), ODD (N=48), and ADHD (N=81). This resulted in a total sample size of 556. Other psychiatric disorders were excluded due to their relatively small sample sizes (less than 20 without comorbidity). For subjects with comorbidity, they were only included once in the group, with the smallest sample-size, to increase statistical power. The disorder symptoms were assessed using a computerized, structured interview (GOASSESS) that was administered to probands, caregivers, or legal guardians, depending on the age of subjects (32). The demographic information for subjects in each group was summarized in Table 1. Data used in this study are publicly available through the database of Genotypes and Phenotypes (dbGaP).

**Table 1.** Demographics information of subjects in the healthy control and disorder groups.

| group | sample size | females | age (mean) | age (std) |
| --- | --- | --- | --- | --- |
| HC | 166 | 82 | 14.51 | 3.79 |
| Specific phobia | 70 | 46 | 12.57 | 3.17 |
| Social phobia | 77 | 43 | 14.05 | 2.85 |
| Depression | 70 | 47 | 16.00 | 2.41 |
| PTSD | 44 | 31 | 15.16 | 2.84 |
| ODD | 48 | 20 | 14.56 | 2.66 |
| ADHD | 81 | 37 | 13.09 | 2.97 |

### MR Image Acquisition

Imaging data were collected in the Hospital of the University of Pennsylvania, using a Siemens Tim Trio (Erlangen, Germany) 3T scanner with 40 mT/m gradients, 200 mT/m/s slew-rates, and a 32-channel head coil optimized for RF transmission. *T*_1_-weighted images were scanned with a 3D, inversion-recovery, magnetization-prepared rapid acquisition gradient echo (MPRAGE) protocol with the following acquisition parameters: TI/TR/TE = 1100/1810/3.51ms, flip angle = 9°, matrix = 256 × 192, FOV= 240 × 180 mm, slices = 160, and slice thickness = 1 mm. DTI images were scanned with a twice-refocused spin-echo single-shot EPI sequence and a custom 64-direction diffusion set, with b-values of 0 and 1000 s/mm^2^. The b = 0 scans were repeated 6 times. Each b = 1000 scan was acquired once at each of the 64 directions, for a total of 70 repetitions. The acquisition parameters for DTI were TR/TE = 8100/82ms, matrix = 128 × 128, FOV= 240 mm, slices = 70, slice thickness = 2 mm and GRAPPA factor = 3. More details of MRI scan protocols and scanner stability for T1-weighted imaging and DTI can be found in a previous study (29).

### Image Preprocessing

We extracted the gray-matter volume (GMV) from the T1-weighted images with the VBM toolbox in SPM8 (Wellcome Trust Centre for Neuroimaging, http://www.fil.ion.ucl.ac.uk/spm/) and fractional anisotropy (FA) from DTI images with PANDA (33). The preprocessing steps included homogeneity bias correction, affine registration, global intensity correction, and segmentation. After segmentation, a Diffeomorphic Anatomic Registration Through Exponentiated Lie algebra algorithm (DARTEL) (34) was run to normalize the segmented images to the MNI-152 space. Then modulation was applied on the normalized images, resulting in the GMV map adjusting for the whole brain size. After smoothing with a 6◻mm full◻width at◻half◻maximum (FWHM) Gaussian kernel, the regional average values of GMV were calculated based on the AAL2 atlas (https://www.gin.cnrs.fr/en/tools/aal/). The processing steps of DTI images included skull removal, correction for simple◻motion and eddy-current distortion, diffusion tensor calculation, and spatial normalization. The FA maps were averaged based on the Johns Hopkins University (JHU) white-matter tractography atlas (35). We chose the tract and label templates which contain 20 and 50 regions, respectively.

### The rationale of multi-dimensional age prediction

In contrast to unidimensional brain-age prediction, we assume that brain development involves multiple subsystems that show staggered developmental pace and distinct patterns of change. If features in different subsystems are mutually exclusive, we can formulate the relationship between chronological age, brain age, and BAG as follows:

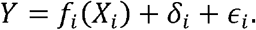

where *X*_*i*_ denotes the brain-imaging features of the *i*^*th*^ subsystem, δ_*i*_ represents the BAG calculated for the *i*^*th*^ subsystem, *Y* is the chronological age, and ϵ_*i*_ is the irreducible error that is assumed to follow a Gaussian distribution with mean 0. Also, imaging features in the *i*^*th*^ sub-system are mapped by function *f*_*i*_ to obtain the brain age of the corresponding subsystem (i.e., *f*_*i*_(*X*_*i*_)).

The multidimensional brain age over *k* different subsystems is denoted as a *k*-dimensional vector (*f*_*1*_(*X*_*1*_), *f*_*2*_(*X*_*2*_),…,*f*_*k*_(*X*_*k*_)). This new metric allows the brain age of some subsystems to be larger than chronological age while the brain age of other subsystems from the same subject are smaller than the chronological age. In contrast, the unidimensional brain-age prediction provides only a single value that summarizes the whole brain’s average developmental status. When we sum the above equation across all *k* subsystems, we have the following:

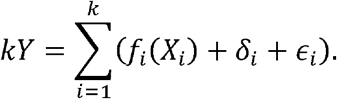

Then,

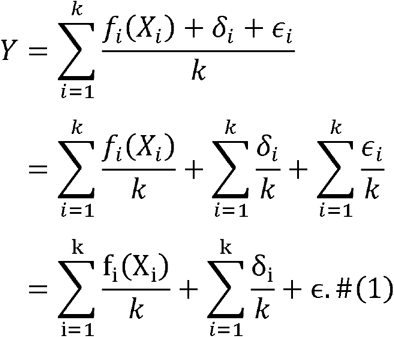

Here ϵ follows a Gaussian distribution with mean 0 as each ϵ_*i*_ follows a Gaussian distribution with mean 0. The first term (i.e., 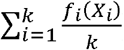 is equivalent to the unidimensional brain age while the second term (i.e., 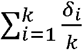) is the BAG under the unidimensional brain age prediction. This illustrates that unidimensional brain age is the average of the multidimensional brain ages across the *k* subsystems.

### Developmental trajectory of brain-imaging features

To identify sub-systems with staggered developmental pace and distinct patterns, we first built a robust regression model to fit each brain-imaging feature using linear and quadratic chronological age, sex, and their interactions as specified in formula (2). In particular, we conducted Huber regression, which is known to be robust to outliers and high leverage points (36).

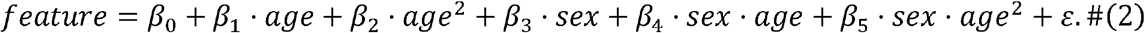

We estimated regression coefficients using the HCs. Given that we had a relatively large sample size and balanced distribution across age, the estimated regression coefficients were used to approximate the developmental trajectory for each brain imaging feature. Other studies have taken a similar approach to approximate the temporal trajectory with cross-sectional data (37–39). To reduce the correlation between the linear and quadratic terms, age was standardized. Imaging features were also standardized before conducting robust regression to make the estimated coefficients comparable across features. Besides, as all the GMV features decreased with age and all the FA features increased with age, we reversed all the FA features. Huber regression parameters were tuned for each feature separately.

We randomly selected 60% of the data as the training set and the remaining 40% as the test set. The tuning parameter that controls the weight of samples with larger residuals was set to be linearly spaced from .1 to 5 with a length of 50. The Huber regression model was fitted to the training data and evaluated using the test data. The *R*^2^ of prediction was computed on the test data. This process was repeated 100 times for each feature with a different seed to split the training and test sets. The optimal tuning parameter was chosen as the one that yielded the highest averaged *R*^2^. To increase the signal-to-noise ratio, the estimated regression coefficients were included in the following analyses only if the *R*^2^ from the corresponding regression model was at least 0.02 (i.e., a small effect size) (40). We did not set a larger threshold to avoid influencing the cluster analyses by removing too many features.

### Clustering of brain-imaging features

We conducted K-means clustering (41) to identify homogeneous subgroups of brain-imaging features based on the estimated coefficients in formula (2)). The data inputted to K-means clustering were in the form of a matrix with each row representing one brain-imaging feature and each column representing one of the regression coefficients in formula (2). Brain-imaging features identified in the same cluster were expected to have a similar developmental trajectory. Cluster analysis has been shown to effectively identify subgroups of functional data or patterns of the curve based on the coefficients of regression (42, 43). We computed the total within-cluster sum of squares (Elbow method) and gap statistics (44) to determine the optimal number of clusters for K-means.

### Multidimensional brain-age prediction

After K-means clustering, we trained the machine-learning model to predict brain age for each cluster using imaging features assigned to this cluster. The predicted brain age from multiple clusters formed a multidimensional brain-age index. The pipeline for the multidimensional brain-age prediction is shown in Fig. 1.

We built a ridge regression model similar to our previous work (19). To build a brain-age prediction model with HCs, a nested 5-fold cross-validation (CV) was conducted. In each CV, 4 folds of the data were assigned to the training set, and the remaining fold was assigned to the test set. An inner 5-fold CV was conducted on the training set to tune the penalty factor of the ridge regression model. Features were scaled based on the training set and then applied to the test set. To predict the brain age of the disorder groups, all HCs were used as the training set to tune the parameter of the ridge regression model with 5-fold CV. The optimized model was then applied to the disorder groups to obtain their brain age estimates. The ridge regression was conducted using the sklearn toolbox v0.22 (45). A unidimensional brain-age prediction was also conducted using all of the imaging features in these clusters according to a similar procedure.

### Bias correction of BAG

After brain-age prediction, we calculated the bias-corrected BAG using the method published in our previous work (19). This correction step removed the negative correlation between the uncorrected BAG and the chronological age (19, 39, 46, 47). This bias correction is essential to reduce confounding when comparing the BAG between groups with different age and sex distributions. As prediction accuracy could be inflated after the bias correction(48), we did not use the bias-corrected BAG for model tuning and comparison.

### Statistical analyses

We evaluated the brain-age prediction performance by the *R*^2^ of a regression model which fitted the brain age with linear and quadratic chronological age, sex, and their interaction effects. Compared to Pearson’s correlation, this method accounts for nonlinear changes in brain age and allows brain age to deviate from the chronological age (19). This was repeated for each cluster and group of subjects. For the HCs, we also examined the correlation between the BAG and chronological age with and without bias-correction for males and females separately. In addition, we compared the corrected BAG between the HCs and each of the disorder groups. We conducted a permutation-based repeated measure ANOVA with the corrected BAG as the dependent variable, the HCs versus disorder group as the between-subject independent variable, and the cluster as the within-subject independent variable. This was repeated for the HCs and each of the six disorder groups separately. For disorder groups that showed significant group by cluster interaction in the permutation-based repeated measure ANOVA, we conducted two-tailed permutation *t*-tests (10,000 permutations) to examine which cluster showed a significant group difference on the corrected BAG. We also conducted permutation *t*-tests to examine whether unidimensional brain age differed significantly between the HCs and the disorder group. The permutation-based approaches allowed us to account for the violation of normality and sphericity assumptions. Lastly, we calculated partial Pearson’s correlation of the corrected BAG calculated from each cluster (i.e., each dimension of the multidimensional brain age) and the unidimensional brain age with sex as a covariate. This was repeated for the HCs and each disorder group separately.

## Results

### Clusters of imaging features with distinct developmental trajectories

Findings from robust regression showed that most brain imaging features followed a clearly nonlinear developmental trajectory with age. Among the 186 features, 184 of them had an *R*^2^ larger than 0.02 (i.e., a small effect size), with a mean *R*^2^ of 0.237. The top GMV features with the highest *R*^2^ were the bilateral superior temporal gyrus and right pallidum. The trajectories of GMV in the bilateral superior temporal gyrus showed similar nonlinear patterns across male and female subjects. In contrast, males on average had a steeper rate of decrease of GMV in the right pallidum than females. The top FA features with the highest *R*^2^ were bilateral cingulum (Fig. 2A). The trajectories of FA features in bilateral cingulum showed distinct patterns between males and females. The distributions of *R*^2^ for all the features were shown in Fig. 2B. On average, GMV features had higher *R*^2^ than FA features, suggesting a larger percentage of variance in GMV can be explained by the estimated developmental trajectory (i.e., robust regression model).

**Figure 2.**
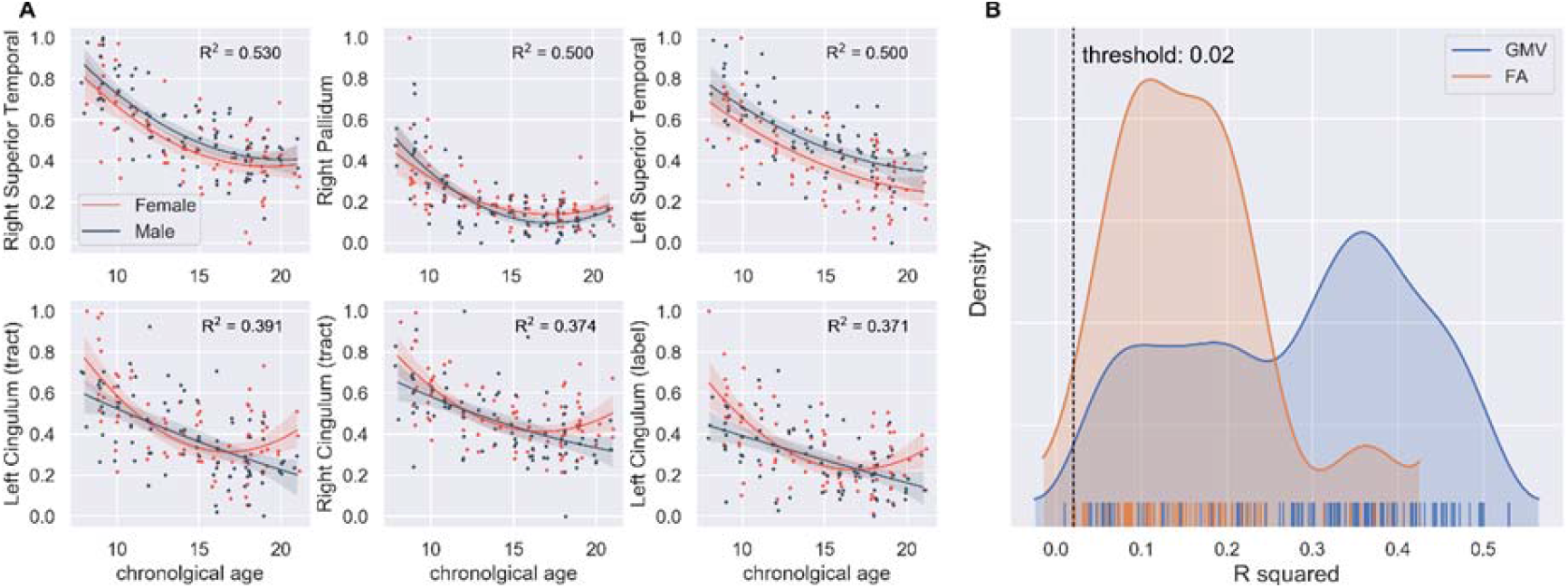
(A) Top 3 GMV and FA features with the best fit in Huber regression. All feature values were scaled from 0 to 1. FA features were reversed before regression to make their change patterns consistent with GMV. The shaded area represented the confidence interval on the 1,000 bootstrap samples. (B) Distribution of *R*^2^ in Huber regression for all GMV and FA features. The *R*^2^ show an obvious mixture of distribution for both GMV and FA. Features with *R*^2^ less than .02 were excluded from later analysis.

Based on the elbow method and gap statistics, we identified that the optimal number of clusters for K-means clustering was four. Most of the GMV and FA features in each cluster showed a symmetric pattern between the left and right hemispheres (Fig. 3). Cluster 1 was composed of features in the cerebellum for both GMV and FA, and in the parahippocampus and temporal pole for GMV. Cluster 2 contained features widely distributed across the frontal lobe, parietal lobe, and the basal ganglia for GMV and in the cingulum and longitudinal fasciculus for FA. The GMV features in cluster 3 were mainly distributed across the frontal orbital and superior occipital regions and the cerebellum. The FA features of cluster 3 were distributed in the corticospinal tract, cerebellar peduncle, as well as internal and external capsule. Cluster 4 contained most of the FA features that were widely distributed across the white-matter tissue. A complete list of brain-imaging features in each cluster is provided in the supplementary materials (Supplementary Table 1).

**Figure 3.**
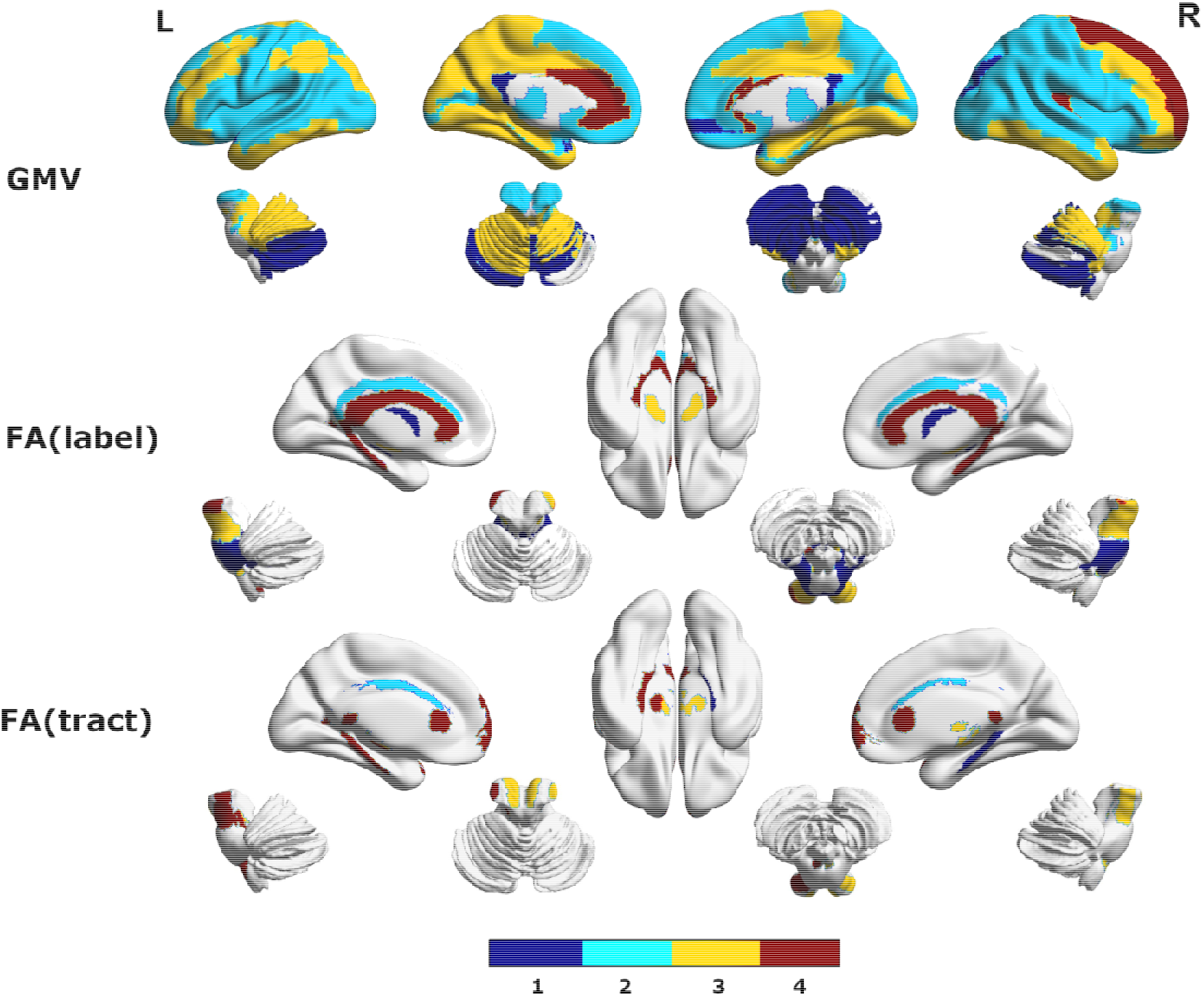
Cluster index for the GMV, FA on label, and tract atlas. In general, the cluster index for most of the regions was symmetrical, which is an indication of the validity of our cluster analysis. Features in cluster 1 were mainly distributed on the cerebellum for both GMV and FA and on the parahippocampus and temporal pole for GMV. Cluster 2 contained the largest number of features that were widely distributed across the frontal, parietal lobe, and the basial ganglia for GMV and on the cingulum and longitudinal fasciculus for FA. Features in cluster 3 were interleaved with those in cluster 2 on the cortical surface. The GMV features were mainly distributed across the frontal orbital region, superior occipital lobe, and the cerebellum. And the FA features were distributed on the corticospinal tract and cerebellar peduncle as well as internal and external capsule. Cluster 4 contains most of the FA features that were widely distributed across the white matter tissue. A complete list of features in each cluster can be found in the supplementary materials (Supplementary Table 1).

After the brain-imaging features were uniquely assigned to one of the four clusters by K-means, we examined the differences across the four clusters. For each cluster, we fitted a regression line to estimate the average trajectory across all the imaging features assigned to this cluster. Additionally, we visualized the aggregated trajectories across all the GMV features and all the FA features separately for each cluster. As illustrated in Supplementary Fig. 1, the aggregated developmental trajectories of brain-imaging features in the four clusters showed distinct patterns. In general, we observed the features in clusters 1 and 4 contained evolutionarily old brain regions such as the limbic system and cerebellum. Features in clusters 2 and 3 contained brain regions in the neocortex that are latterly evolved. The small linear trend of features in clusters 1 and 4 indicates these regions develop at the early stage of life, which is not covered in our samples. In addition, clusters 2 and 3 contained regions in most of the established networks such as the default mode network (49), dorsal attention network (50), and sensory-motor network (51).

As clustering was performed on the estimated regression coefficients, we computed the average coefficient values across the brain-imaging features assigned to each cluster (Table 2). In addition, we generated scatter and density plots of the regression coefficients for brain-imaging features in each cluster (Fig. 4). As shown in Table 2, features in cluster 2 had a relatively large coefficient of age (*b* = −0.414) while features in cluster 4 had a relatively large, averaged coefficient of age^2^ (*b* = 0.143). In addition, features in cluster 1 had a relatively large interaction effect between sex and the linear term of age (*b* = 0.051) while features in cluster 4 had a relatively large interaction effect between sex and the quadratic term of age (*b* = −0.108). This suggests that different clusters demonstrated different developmental trajectories. In the subpanel of Fig. 4 with intercept as the y-axis and the coefficient of age^2^ as the x-axis (highlighted subpanel 1), we see a clear separation between clusters 1 and 4 along the diagonal line. Additionally, in the subpanel with the coefficient of age as the y-axis and the coefficient of sex by age^2^ as the x-axis (highlighted subpanel 2), we observe a better separation between the four clusters.

**Table 2.** Regression coefficients averaged across brain imaging features in each cluster.

| Cluster | Intercept | Age | Age<sup>2</sup> | Sex | Sex*Age | Sex*Age<sup>2</sup> | Cluster<br>Size |
| --- | --- | --- | --- | --- | --- | --- | --- |
| 1 | 0.019 | -0.106 | 0.003 | 0.041 | 0.051 | -0.044 | 29 |
| 2 | -0.067 | -0.414 | 0.115 | 0.086 | -0.038 | -0.003 | 68 |
| 3 | -0.053 | -0.299 | 0.074 | -0.044 | -0.046 | -0.008 | 46 |
| 4 | -0.117 | -0.193 | 0.143 | 0.098 | -0.024 | -0.108 | 41 |

**Figure 4.**
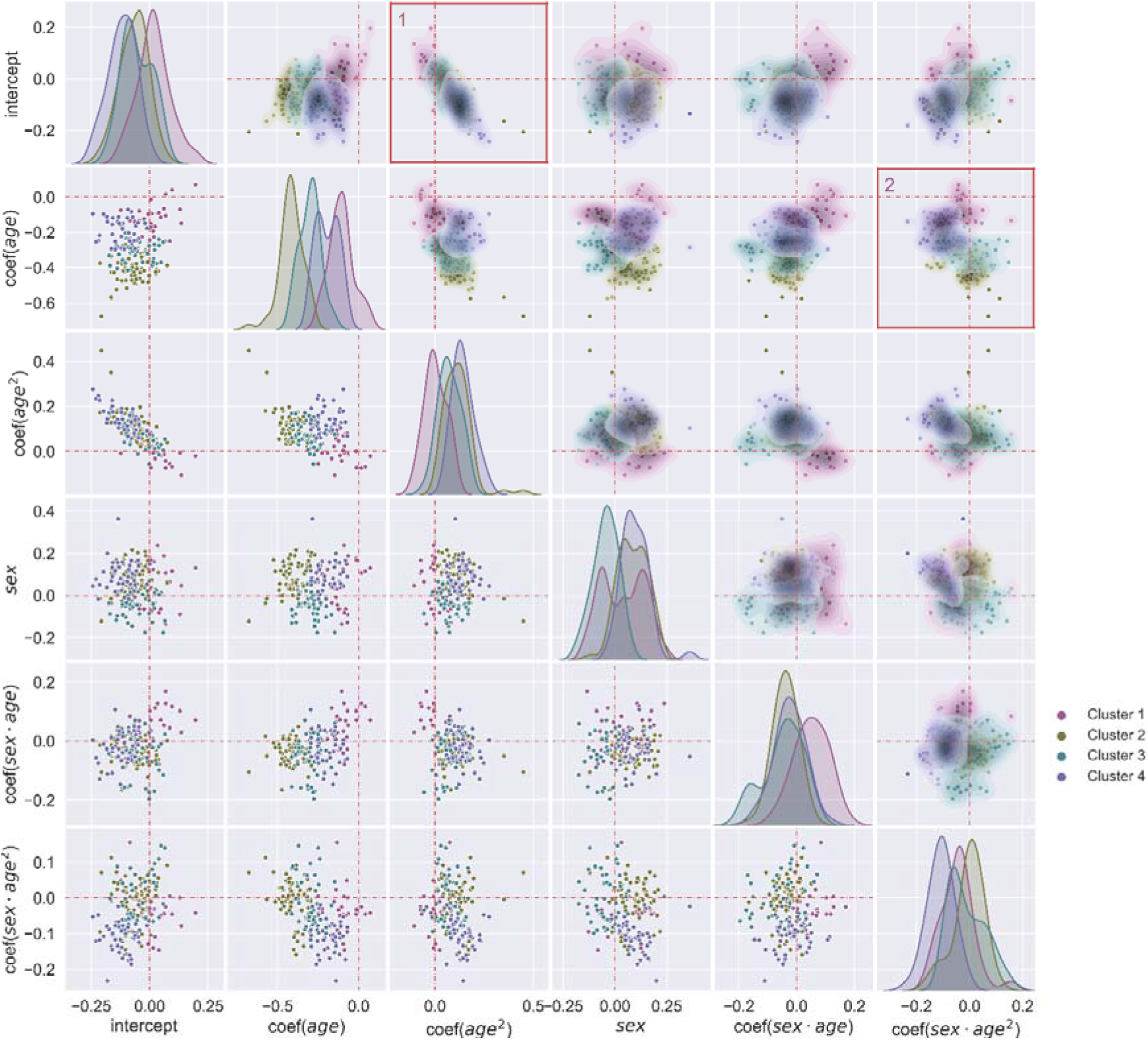
Scatter and density plots of the coefficients in Huber regression for brain-imaging features in each cluster. Features in cluster 1 had small coefficients for the linear and quadratic terms. Features in cluster 2 had large coefficients for the linear quadratic terms. Features in cluster 3 had large coefficients for the linear term and moderate coefficients for the quadratic term. Features in cluster 4 had small coefficients for the linear term and large coefficients for the quadratic term. Additionally, features in cluster 4 had a relatively large interaction effect between sex and the quadratic term of age while features in cluster 1 had a relatively large interaction effect between sex and the linear term of age. Features in cluster 2, 3, and 4 had a moderate interaction effect between sex and the linear term of age. This effect was opposite to that of cluster 1.

### Multidimensional brain-age index

After K-means clustering, we conducted brain age prediction for each cluster separately. The predicted brain age from each of the four clusters formed a four-dimensional brain-age index. In addition, we computed the brain-age prediction using all imaging features from these clusters, which yielded the unidimensional brain-age index. The trajectories of brain age, BAG, and corrected BAG were visualized in Fig. 5. Each of the four clusters showed a distinct developmental trajectory of the brain age (first row in Fig. 5). The developmental trajectory was relatively linear in cluster 1 while the remaining clusters showed more nonlinear patterns. This was consistent with their averaged coefficients in Table 2 as cluster 1 had the smallest coefficient of age^2^. For cluster 2 and 3, females had a slightly larger brain age than males during early adolescence. This trend reversed during middle adolescence for cluster 3 and at early adulthood for cluster 2. For cluster 4, the trajectory of brain age for females was nonlinear while the trajectory for males was approximately linear over time. For cluster 1, females and males had very similar brain age during middle adolescence. However, females had lower brain age than males in early and late adolescence. In addition, cluster 2 achieved the highest prediction accuracy (*R*^2^ = .735 for males and *R*^2^ = .659 for females), similar to the prediction accuracy with all imaging features (*R*^2^ = .741 for males and *R*^2^ = .650 for females). The prediction accuracy was the lowest in cluster 1 (*R*^2^ = .173 for males and *R*^2^ = .209 for females). This might be explained by the observation that cluster 2 included a larger number of imaging features than cluster 1 (Table 2). In addition, we also observed a consistent negative correlation between the BAG and the chronological age. This bias was smaller for clusters with higher prediction accuracy, as indicated by smaller correlation coefficients (second row in Fig. 5). Lastly, our bias-correction method successfully removed the negative correlation (third row in Fig. 5).

**Figure 5.**
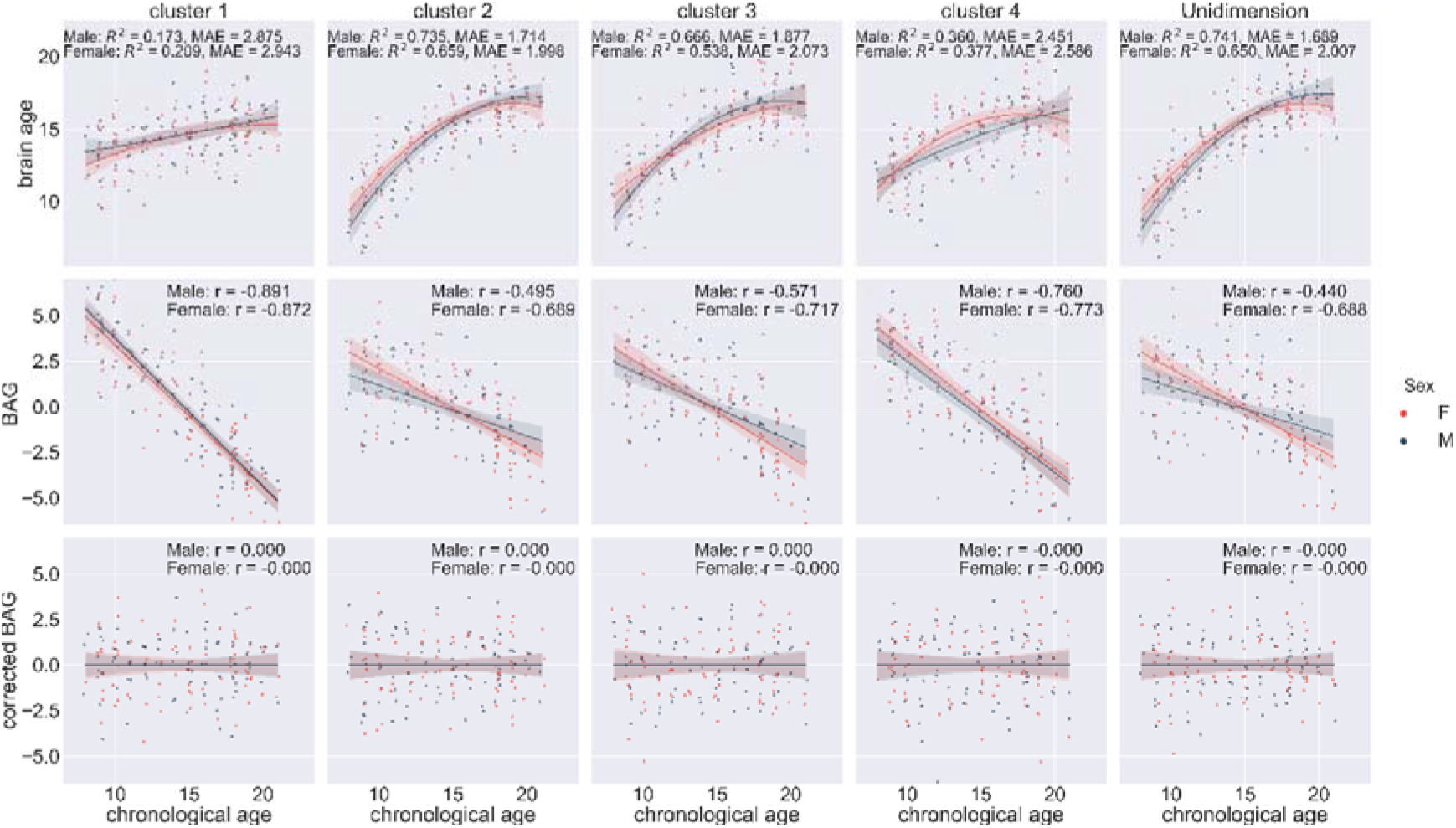
Brain-age prediction and bias-correction using features in each cluster and all clusters combined (unidimensional brain age). The first row showed the predicted age (brain age) versus the chronological age for each cluster and unidimensional brain-age prediction. The second row showed the corresponding BAG (brain age – chronological age) versus the chronological age. The third row showed the corresponding BAG with bias-correction versus the chronological age. The shaded area represented the confidence interval on the 1,000 bootstrap samples.

### Cluster-specific alterations of brain development in psychiatric disorders

To examine whether the bias-corrected BAG obtained from different clusters demonstrated differential ability in separating disorder groups from HCs, we ran a permutation-based repeated measure ANOVA with a two-level factor group (disorder group vs. HCs), a four-level factor cluster (each of the four clusters identified in K-means), and their interaction. As shown in Table 3, we found a significant main effect of cluster for specific phobia (*p* = .001), with FDR correction for multiple comparisons. In addition, the interaction effect was significant for specific phobia (*p* =.001), depression (*p* = .011), and ADHD (*p* = .004), with FDR correction for multiple comparison. All these effects remained significant even with a more conservative Holm correction.

**Table 3.** Main and interaction effects of mixed ANOVA on the corrected BAG (with FDR correction for multiple comparisons) between each of the disorder groups and HCs.

| Disorder | *p* (Group) | *p* (Cluster) | *p* (Interaction) |
| --- | --- | --- | --- |
| Specific phobia | 0.373 | 0.001** | 0.001** |
| Social phobia | 0.975 | 0.904 | 0.905 |
| Depression | 0.975 | 0.005** | 0.011* |
| PTSD | 0.975 | 0.069 | 0.064 |
| ODD | 0.211 | 0.772 | 0.777 |
| ADHD | 0.975 | 0.004** | 0.004** |

To better understand the interaction between Group and Cluster, we conducted permutation *t*-tests on the corrected BAG between the disorder groups and HCs that showed significant interaction effects for each cluster separately (Table 4 and Fig. 6, A). Compared to HCs, the corrected BAG of the specific phobia group was significantly smaller in cluster 1 (Cohen’s *d*◻=◻−.459, *p* = .012), cluster 3 (Cohen’s *d*◻=◻−.361, *p* = .049), and for the unidimensional brain-age gap (*p* = .049), with FDR correction for multiple comparisons. This finding is consistent with the significant interaction effect identified from repeated measure ANOVA for specific phobia. It also suggests that different clusters showed differential ability to separate the specific phobia group from HCs in terms of their corrected BAG. In addition, the unidimensional brain age was only significantly associated with specific phobia (Cohen’s *d*◻=◻−.367, *p* = .049) while the multidimensional brain age index showed an additional significant difference in cluster 1 for ADHD (Cohen’s *d*◻=◻−.352, *p* = .049) and marginally significant difference in cluster 4 for depression (Cohen’s *d*◻=◻−.370, *p* = .060).

**Table 4.** Comparison of mean difference of the corrected BAG between each disorder group and HCs with permutation t-test with FDR correction.

|  | Cluster 1 |  | Cluster 2 |  | Cluster 3 |  | Cluster 4 |  | Unidimension |  |
| --- | --- | --- | --- | --- | --- | --- | --- | --- | --- | --- |
|  | *Cohen's d* | *p* | *Cohen's d* | *p* | *Cohen's d* | *p* | *Cohen's d* | *p* | *Cohen's d* | *p* |
| Specific phobia | -0.459 | 0.012* | -0.144 | 0.512 | -0.361 | 0.049* | 0.304 | 0.114 | -0.367 | 0.049* |
| Depression | 0.171 | 0.417 | -0.032 | 0.836 | 0.099 | 0.773 | -0.370 | 0.060 | -0.057 | 0.812 |
| ADHD | -0.352 | 0.049* | 0.029 | 0.836 | -0.066 | 0.792 | 0.180 | 0.417 | -0.073 | 0.792 |

**Figure 6.**
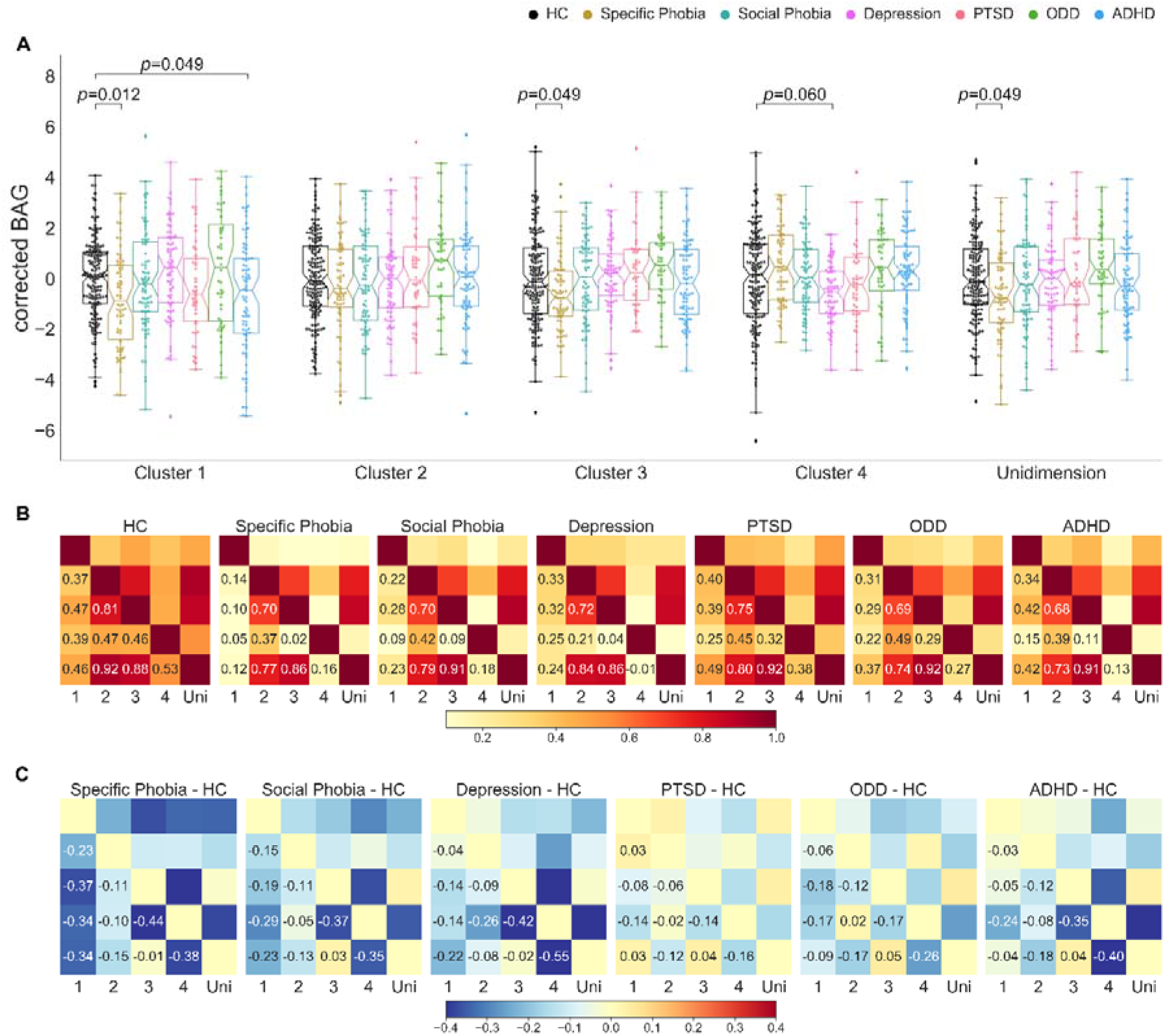
A) Permutation *t*-test between the brain-age gap (with bias-correction, BAG) for HCs and each of the disorder groups. The bias of BAG was corrected by the model that residualized the linear and quadratic trend of the age, sex, and their interactions. (A) The corrected BAGs were significantly lower than the HCs for specific phobia on cluster 1 (Cohen’s *d*◻=◻-.459, *p* = .012), cluster 3 (Cohen’s *d*◻=◻-.361, *p* = .049), and unidimensional BAG (Cohen’s *d*◻=◻-.367, *p* = .049), ADHD on cluster 1 (Cohen’s *d*◻=◻−.352, *p* = .049), and marginally significant for depression (Cohen’s *d*◻=◻-.370, *p* = .060) with FDR correction for multiple comparison. (B) Similarity matrix of Pearson's correlation between the corrected BAG obtained from each pair of clusters for HCs and each disorder group. On the HCs, the BAG of the unidimensional brain age was highly correlated with that of the cluster 2 (*r* = .92) and 3 (*r* = .88), and moderately correlated with cluster 1 (*r* = .46) and 4 (*r* = .53). The BAGs of cluster 2 and 3 are highly correlated (*r* = .81). The BAG between cluster 1 and 3 (*r* = .47), cluster 2 and 4 (*r* = .47), cluster 3 and 4 (*r* = .46) were moderately correlated. The BAG correlations between cluster 1 and 2 (*r* = .37) and between cluster 1 and 4 (*r* = .39) were relatively weaker than the correlations between other clusters. (C) Difference between the correlation matrix of each disorder group and HCs. For most of the pairs of correlations, values in the correlation matrix for the disorder groups are lower than those of the HCs, especially for the pairs between cluster 1 and others and the pairs between cluster 3 and 4.

To examine whether the corrected BAG obtained from different clusters contained complementary information on precocious or delayed development of different brain systems, we calculated partial Pearson’s correlation of the corrected BAG between each pair of clusters for the HCs and disorder groups separately with sex as a covariate. The correlation matrix was illustrated as a similarity of developmental patterns across brain subsystems (clusters). Among HCs, clusters 2 and 3 were highly corrected (*r* = .81) in terms of the corrected BAG (Fig. 6, B). The unidimensional brain age was highly correlated with the corrected BAG calculated cluster 2 (*r* = .92) and cluster 3 (*r* = .88). Other correlations were considered moderate. In general, the correlations calculated for the psychiatric disorder groups were weaker than what we observed for HCs. This suggests that alterations in brain development caused by disorders varied across subjects and brain systems due to their heterogeneous characteristics. To further illustrate this point, we subtracted the correlation matrix of HCs from the correlation matrix of each of the disorder groups (Fig. 6, C). In general, we observed that the difference in the correlations was negative. In particular, for specific phobia, the correlations of the corrected BAG between cluster 1 and other clusters were largely reduced compared to those for the HCs. We also observed a large reduction in the correlation of the corrected BAG between clusters 3 and 4 for specific phobia, social phobia, depression, and ADHD. For PTSD and ODD, the reduction in correlation was moderate for most of the cluster pairs.

Additionally, we examined how multimodal brain-age prediction revealed altered brain-age developmental trajectory in the disorder groups that were otherwise invisible to unidimensional brain-age prediction. As illustrated in Fig. 7, the brain-age prediction with the 4 clusters showed altered brain developmental trajectories for the disorder groups. Specifically, compared to the HCs, brain age obtained with cluster 1 of the disorder groups was lower in early adolescence but returned to normal subsequently. In addition, for depression and ODD, the brain developmental trajectory had a strong nonlinear trend for cluster 1. This reflected a higher quadratic tend of features in the disorder groups than the HCs which may indicate an earlier deterioration of this brain system. For clusters 2 and 3, the depression groups showed lower brain age in early adolescence, whereas the PTSD group showed higher brain age in this stage. The brain age in cluster 2 of the ODD group showed a strong nonlinear trend similar to what was observed in cluster 1, with higher brain age in middle adolescence and lower brain age in late adolescence. For cluster 4, all the disorder groups showed higher brain age in early adolescence and lower brain age in late adolescence. Importantly, all of these alterations were attenuated in the unidimensional brain-age predictions.

**Figure 7.**
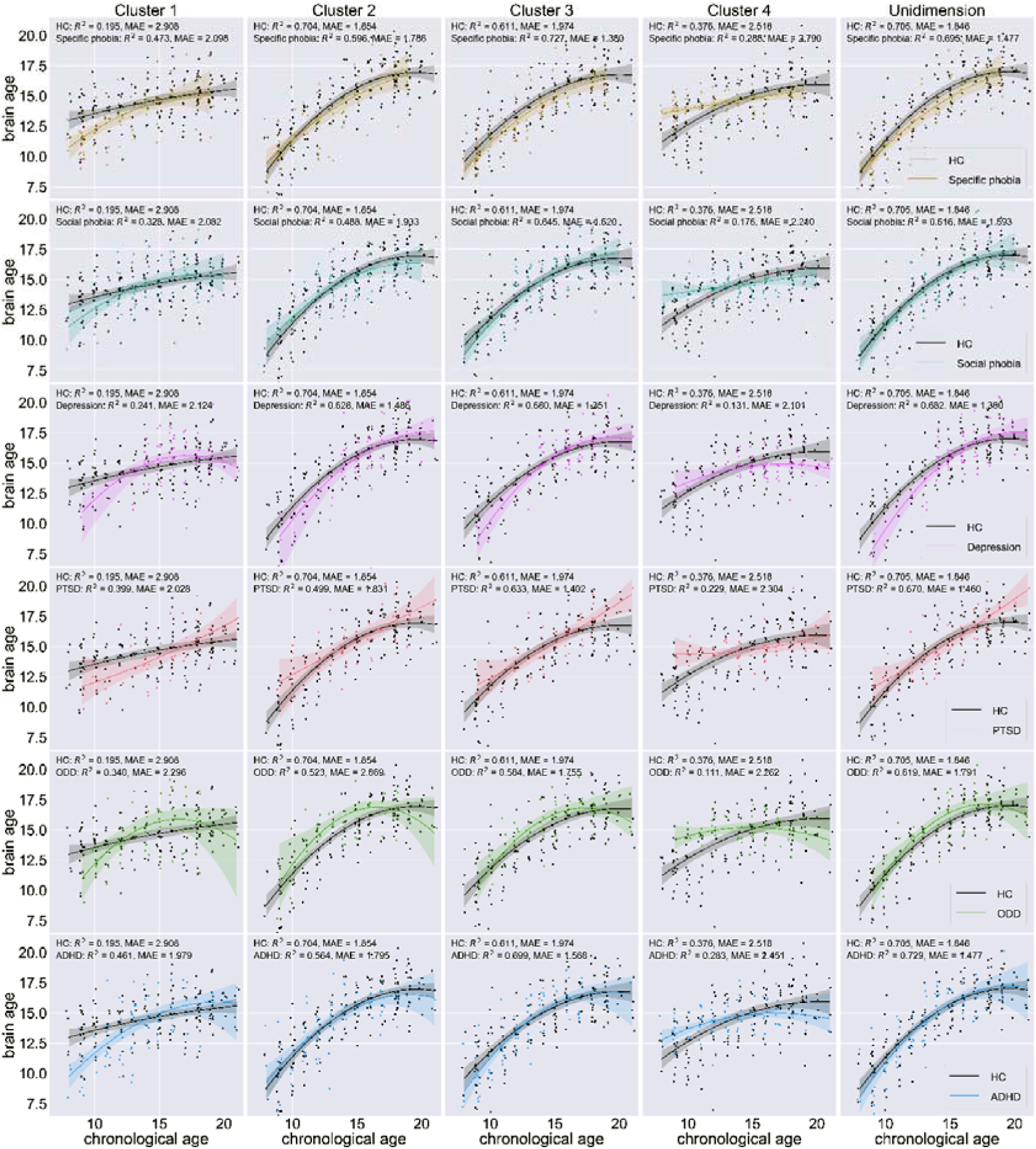
Altered brain development trajectories on the disorder groups revealed by multimodal brain-age prediction. The scatterplots showed the predicted age (brain age) versus the chronological age for the HCs and each of the disorder groups obtained from brain-imaging features in each cluster and all clusters combined. The shaded area represents the confidence interval on 1,000 bootstrap samples.

## Discussion

We classified multimodal brain-imaging features into different clusters based on their developmental trajectories delineated by the coefficients in a robust regression model. For each cluster, we obtained brain ages using a ridge regression model. The change in brain age for different clusters showed distinct trajectories in accordance with the corresponding features in each cluster. We also found that the BAGs derived from our proposed MBAI approach captured abnormalities that were constrained in subsystems of the brain and which were not detectable to unidimensional brain age prediction. Finally, the BAGs of different clusters showed moderate correlations with each other in the HCs and reduced correlations in the disorder groups. This indicated that the BAGs from different clusters contained unique information about brain development which holds the potential to serve as novel biomarkers for psychiatric disorders.

### The nonlinearity of brain developmental trajectory

The linearity of brain age prediction was mixed in previous studies. Some studies showed equally accurate age prediction across the lifespan (23, 24, 27), whereas others showed clearly nonlinear brain-age patterns (19, 20, 47). Our results showed divergent developmental patterns of brain age. Specifically, features in clusters 1 and 2 showed moderate coefficients for the quadratic term age, whereas those in clusters 3 and 4 showed low coefficients. The brain ages obtained for each cluster were consistent with the developmental patterns of features.

Theoretically, we postulate a nonlinear brain developmental trajectory across a wide lifespan. However, as different brain structures show rapid development at different stages in lifespan, a unitary brain-age prediction model may have varying weights on brain-imaging features at different age ranges to achieve an accurate prediction for the chronological age. This is especially prominent for ML models with nonlinear kernels or deep neural networks with complex structures (4, 23, 27, 52). It is worth noting that features that did not show obvious quadratic developmental patterns in the current study also did not show a large linear trend. This implies that the age range of rapid developmental changes in these features is not covered by our study. There is evidence showing that the most drastic changes in the brain take place during the first 5 years of life (53). By using a sample with a wider age range, it is likely that we can better categorize features with distinct developmental patterns.

### Divergent developmental trajectories of different brain systems

As brain structure shows divergent developmental trajectories for different regions (12, 54), we proposed a multidimensional brain-age prediction model that is designed to capture distinct brain developmental trajectories. Recent research has suggested that modeling distinct and regional patterns of aging may yield additional biomarkers for health and disease. One study examining altered maternal brain development on subjects between 45-82 years conducted hierarchical clustering on the MRI features (38). Another study used independent component analysis (ICA) to decompose the brain imaging features into 62 modes of subject variability (14). The ICA was also directly conducted on the original imaging features. Unlike these studies (38, 14) and other ones that conducted separate brain age prediction based on imaging modality (24, 55), we first ran robust regression to extract the coefficients that contained information of developmental pattern of each feature. Then, K-means cluster analysis was conducted based on the coefficients to group features into clusters with different developmental patterns. In this sense, our proposed method is better at capturing distinct brain developmental trajectories by clustering imaging features based on their coefficients estimated from robust regression. Compared to other popular clustering methods such as hierarchical and Gaussian mixture model clustering, the K-means algorithm is fast and yields more balanced groups (56, 57). Even though we do not assume that the features are balanced across clusters, a small number of features in one cluster may lead to poor brain-age prediction performance. A systematic examination of different clustering algorithms for multi-dimensional brain-age prediction is still needed in future work with larger sample sizes, wider age ranges through the lifespan, and more refined brain parcellations (58).

### More Sensitive brain age-based analyses

As we assume the existence of BAG in the age prediction model, a good predictive model for brain-age estimation should not overfit the data and yield perfect prediction for chronological age. Typical ways to avoid overfitting include imposing a higher level of regularization (27) and using lower-dimensional linear models instead of models with nonlinear kernels. In this work, we chose ridge regression rather than complex models with nonlinear kernels, which can result in perfect prediction for the chronological age but tend to overfit for brain-age prediction. In addition, our approach utilizing a reduced number of features in each cluster mitigates the “curse of dimensionality” and thus is more sensitive to brain developmental alterations. Unlike sparse models such as the LASSO and elastic net (59, 60) which only select the most predictive features, our method preserves features that are less predictive for chronological age but still contains information about the precocity and delay of brain development. As different brain regions have complex functional interactions, a more sophisticated decomposition method to identify the multidimensional structure of brain development should be the subject of future work.

### Altered brain development in psychiatric disorders

Our results revealed that psychiatric disorders were related to altered brain development of distinct and overlapping brain systems. Specifically, for the averaged BAG across the entire adolescence stage, a smaller BAG was found in specific phobia for features in clusters 1 and 3 and ADHD for cluster 1. The depression group also showed a lower BAG with marginal significance for cluster 4. Taking different stages of adolescence into consideration, a smaller BAG for cluster 1 and a larger BAG for cluster 4 were found for most of the disorders during early adolescence. Previous studies found that specific phobia was associated with alterations in the amygdala, insula, anterior cingulate cortex, dorsolateral prefrontal cortex (61, 62), and FA in inferior longitudinal fasciculus, inferior fronto-occipital fasciculus, and uncinate fasciculus (63).

ADHD with frequent comorbidity on ODD was associated with alterations in GMV of orbitofrontal cortex, the superior frontal gyrus, middle frontal gyrus, and the inferior parietal gyrus (64) and FA of inferior fronto-occipital fasciculus, corticospinal tract, forceps minor, and internal capsule (65). In addition, ADHD is closely related to the dysfunction of frontal–striatal– cerebellar circuits (66). Some of these regions are consistent with features in the altered cluster-based age prediction in the disorder groups. For example, the GMV of the amygdala, anterior cingulate cortex, orbitofrontal, and corticospinal tract that showed abnormalities in specific phobia were assigned to cluster 3. The GMV of a wide range of cerebellum regions were assigned to cluster 1. The GMV of the superior frontal gyrus and FA of the superior fronto-occipital fasciculus were assigned to cluster 4. In summary, the multidimensional brain-age prediction revealed brain alterations in constrained regions that would otherwise be invisible to unidimensional age prediction.

In addition to examining the BAG separately for each dimension, we proposed to examine the pattern of developmental similarity across brain systems using partial correlations between pairs of multidimensional BAGs with sex as a covariate. We found moderate correlations between the developmental indices across brain systems. This implies that brain development is determined by both a general factor that influences the overall development and domain-specific factors that influence one or more of the brain subsystems. Furthermore, we showed reduced developmental similarity patterns in the disorder groups, which was potentially caused by differential alterations of the disorder across subjects with different chronological ages. This implies that alterations of brain structure in psychiatric disorders could be moderated by chronological age. The moderating effect of chronological age, which varies across brain systems, can be modeled with Gaussian processes that characterize the developmental fingerprints for psychiatric disorders. With longitudinal data, the similarity of developmental alterations across different brain systems can be used as a biomarker for the diagnosis of psychiatric disorders and a guide for early interventions for vulnerable children.

### Future directions

There are several important issues in brain-age prediction yet to be explored. First, brain development is characterized by both structural and functional changes (12). It has been shown that resting-state functional connectivity yields comparable age prediction performance (24), but how it is related to psychiatric disorders has not been examined. Besides, how brain activity during basic psychological tasks is related to brain development is yet to be examined. Second, higher-order brain-imaging metrics such as small-worldness (67) and controllability (68) of the brain network are also related to brain development and cognition and likely provide additional information for brain-age prediction. Finally, although we assumed static and non-overlapping brain-structure clusters in the current study, the development of different aspects of cognition and the progression of psychiatric disorders involve both distinct and shared brain systems and manifest as dynamic processes dependent on life stages, which may be examined in future research.

## Acknowledgments

We would like to thank Mary V. Spiers for helpful discussion on this project, and the reviewers for their thorough work.

## References

1. F. M. Benes, M. Turtle, Y. Khan, P. Farol, Myelination of a key relay zone in the hippocampal formation occurs in the human brain during childhood, adolescence, and adulthood. Arch. Gen. Psychiatry 51, 477–84 (1994).

2. D. Purves, J. W. Lichtman, Elimination of synapses in the developing nervous system. Science 210, 153–7 (1980).

3. N. Gogtay, et al., Dynamic mapping of human cortical development during childhood through early adulthood. Proc. Natl. Acad. Sci. 101, 8174–8179 (2004).

4. H. C. Hazlett, et al., Early brain development in infants at high risk for autism spectrum disorder. Nature 542, 348–351 (2017).

5. A. Giorgio, et al., Changes in white matter microstructure during adolescence. Neuroimage 39, 52–61 (2008).

6. E. L. Dennis, et al., Development of brain structural connectivity between ages 12 and 30: A 4-Tesla diffusion imaging study in 439 adolescents and adults. Neuroimage 64, 671–684 (2013).

7. T. R. Insel, B. N. Cuthbert, Brain disorders? Precisely: Precision medicine comes to psychiatry. Science (80-.). 348, 499–500 (2015).

8. D. C. Hesdorffer, Comorbidity between neurological illness and psychiatric disorders. CNS Spectr. 21, 230–238 (2016).

9. A. T. Drysdale, et al., Resting-state connectivity biomarkers define neurophysiological subtypes of depression. Nat. Med. 23, 28–38 (2017).

10. J. N. Giedd, et al., Brain development during childhood and adolescence: a longitudinal MRI study. Nat. Neurosci. 2, 861–863 (1999).

11. T. Paus, M. Keshavan, J. N. Giedd, Why do many psychiatric disorders emerge during adolescence? Nat. Rev. Neurosci. 9, 947–957 (2008).

12. T. Paus, Mapping brain maturation and cognitive development during adolescence. Trends Cogn. Sci. 9, 60–68 (2005).

13. B. A. Jonsson, et al., Brain age prediction using deep learning uncovers associated sequence variants. Nat. Commun. 10 (2019).

14. S. M. Smith, et al., Brain aging comprises many modes of structural and functional change with distinct genetic and biophysical associations. Elife 9 (2020).

15. K. Franke, C. Gaser, Longitudinal Changes in Individual BrainAGE in Healthy Aging, Mild Cognitive Impairment, and Alzheimer’s Disease. GeroPsych (Bern). 25, 235–245 (2012).

16. N. Koutsouleris, et al., Accelerated brain aging in schizophrenia and beyond: a neuroanatomical marker of psychiatric disorders. Schizophr. Bull. 40, 1140–53 (2014).

17. Y. Chung, et al., Use of Machine Learning to Determine Deviance in Neuroanatomical Maturity Associated With Future Psychosis in Youths at Clinically High Risk. JAMA Psychiatry (2018) https:/doi.org/10.1001/jamapsychiatry.2018.1543 (July 16, 2018).

18. J. H. Cole, K. Franke, Predicting Age Using Neuroimaging: Innovative Brain Ageing Biomarkers. Trends Neurosci. 40, 681–690 (2017).

19. X. Niu, F. Zhang, J. Kounios, H. Liang, Improved prediction of brain age using multimodal neuroimaging data. Hum. Brain Mapp., hbm.24899 (2019).

20. N. U. F. Dosenbach, et al., Prediction of individual brain maturity using fMRI. Science 329, 1358–61 (2010).

21. B. Mwangi, K. M. Hasan, J. C. Soares, Prediction of individual subject’s age across the human lifespan using diffusion tensor imaging: A machine learning approach. Neuroimage 75, 58–67 (2013).

22. G. Erus, et al., Imaging Patterns of Brain Development and their Relationship to Cognition. Cereb. Cortex 25, 1676–1684 (2015).

23. J. H. Cole, et al., Predicting brain age with deep learning from raw imaging data results in a reliable and heritable biomarker. Neuroimage 163, 115–124 (2017).

24. M. Truelove-Hill, et al., A multidimensional Neural Maturation Index reveals reproducible developmental patterns in children and adolescents. J. Neurosci. 40, 2092–19 (2020).

25. H. M. Aycheh, et al., Biological brain age prediction using cortical thickness data: A large scale cohort study. Front. Aging Neurosci. 10, 252 (2018).

26. F. Liem, et al., Predicting brain-age from multimodal imaging data captures cognitive impairment. Neuroimage 148, 179–188 (2017).

27. V. M. Bashyam, et al., MRI signatures of brain age and disease over the lifespan based on a deep brain network and 14◻468 individuals worldwide. Brain 143, 2312–2324 (2020).

28. R. C. Gur, et al., A cognitive neuroscience-based computerized battery for efficient measurement of individual differences: Standardization and initial construct validation. J. Neurosci. Methods 187, 254–262 (2010).

29. T. D. Satterthwaite, et al., Neuroimaging of the Philadelphia Neurodevelopmental Cohort. Neuroimage 86, 544–553 (2014).

30. T. D. Satterthwaite, et al., Neuroimaging of the Philadelphia Neurodevelopmental Cohort. Neuroimage 86, 544–553 (2014).

31. R. C. Gur, et al., Age group and sex differences in performance on a computerized neurocognitive battery in children age 8−21. Neuropsychology 26, 251–265 (2012).

32. M. E. Calkins, et al., The Philadelphia Neurodevelopmental Cohort: constructing a deep phenotyping collaborative. J. Child Psychol. Psychiatry 56, 1356–1369 (2015).

33. Z. Cui, S. Zhong, P. Xu, Y. He, G. Gong, PANDA: a pipeline toolbox for analyzing brain diffusion images. Front. Hum. Neurosci. 7, 42 (2013).

34. J. Ashburner, A fast diffeomorphic image registration algorithm. Neuroimage (2007) https:/doi.org/10.1016/j.neuroimage.2007.07.007 (November 3, 2020).

35. K. Hua, et al., Tract probability maps in stereotaxic spaces: Analyses of white matter anatomy and tract-specific quantification. Neuroimage 39, 336–347 (2008).

36. J. Fox, S. Weisberg, Robust regression. An R S-Plus companion to Appl. Regres. 91 (2002).

37. T. T. Brown, et al., Neuroanatomical Assessment of Biological Maturity. Curr. Biol. 22, 1693–1698 (2012).

38. A. G. Lange, et al., The maternal brain: Region◻specific patterns of brain aging are traceable decades after childbirth. Hum. Brain Mapp., hbm.25152 (2020).

39. S. M. Smith, D. Vidaurre, F. Alfaro-Almagro, T. E. Nichols, K. L. Miller, Estimation of brain age delta from brain imaging. Neuroimage 200, 528–539 (2019).

40. J. Cohen, Statistical power analysis for the behavioral sciences. Lawrence Earlbam Assoc. Hillsdale, NJ (1988).

41. S. P. Lloyd, Least Squares Quantization in PCM. IEEE Trans. Inf. Theory 28, 129–137 (1982).

42. T. Tarpey, Linear transformations and the k-means clustering algorithm: Applications to clustering curves. Am. Stat. 61, 34–40 (2007).

43. J. Jacques, C. Preda, Functional data clustering: A survey. Adv. Data Anal. Classif. 8, 231–255 (2014).

44. R. Tibshirani, G. Walther, T. Hastie, Estimating the number of clusters in a data set via the gap statistic. J. R. Stat. Soc. Ser. B Stat. Methodol. 63, 411–423 (2001).

45. F. Pedregosa, et al., “Scikit-learn: Machine Learning in Python” (2011).

46. H. Liang, F. Zhang, X. Niu, Investigating systematic bias in brain age estimation with application to post◻traumatic stress disorders. Hum. Brain Mapp., hbm.24588 (2019).

47. T. T. Le, et al., A Nonlinear Simulation Framework Supports Adjusting for Age When Analyzing BrainAGE. Front. Aging Neurosci. 10, 317 (2018).

48. E. Butler, et al., Statistical Pitfalls in Brain Age Analyses. bioRxiv, 2020.06.21.163741 (2020).

49. M. D. Fox, M. E. Raichle, Spontaneous fluctuations in brain activity observed with functional magnetic resonance imaging. Nat. Rev. Neurosci. 8, 700–711 (2007).

50. M. D. Fox, et al., The human brain is intrinsically organized into dynamic, anticorrelated functional networks. Proc. Natl. Acad. Sci. U. S. A. 102, 9673–9678 (2005).

51. A. Londei, et al., Sensory-motor brain network connectivity for speech comprehension. Hum. Brain Mapp. 31, NA-NA (2009).

52. K. Franke, G. Ziegler, S. Klöppel, C. Gaser, Estimating the age of healthy subjects from T1-weighted MRI scans using kernel methods: Exploring the influence of various parameters. Neuroimage 50, 883–892 (2010).

53. J. Levman, P. MacDonald, A. R. Lim, C. Forgeron, E. Takahashi, A pediatric structural MRI analysis of healthy brain development from newborns to young adults. Hum. Brain Mapp. 38, 5931–5942 (2017).

54. R. K. Lenroot, J. N. Giedd, Brain development in children and adolescents: Insights from anatomical magnetic resonance imaging. Neurosci. Biobehav. Rev. 30, 718–729 (2006).

55. G. Hwang, et al., Brain aging in temporal lobe epilepsy: Chronological, structural, and functional. NeuroImage Clin. 25 (2020).

56. R. Xu, D. C. Wunsch, Clustering algorithms in biomedical research: A review. IEEE Rev. Biomed. Eng. 3, 120–154 (2010).

57. J. A. Hartigan, M. A. Wong, Algorithm AS 136: A K-Means Clustering Algorithm. Appl. Stat. 28, 100 (1979).

58. C. Wiwie, J. Baumbach, R. Röttger, Comparing the performance of biomedical clustering methods. Nat. Methods 12, 1033 (2015).

59. H. Zou, T. Hastie, Regularization and variable selection via the elastic net. J. R. Stat. Soc. Ser. B (Statistical Methodol. 67, 301–320 (2005).

60. R. Tibshirani, Regression Shrinkage and Selection Via the Lasso. J. R. Stat. Soc. Ser. B 58, 267–288 (1996).

61. J. C. Ipser, L. Singh, D. J. Stein, Meta-analysis of functional brain imaging in specific phobia. Psychiatry Clin. Neurosci. 67, 311–322 (2013).

62. C. I. Wright, B. Martis, K. McMullin, L. M. Shin, S. L. Rauch, Amygdala and insular responses to emotionally valenced human faces in small animal specific phobia. Biol. Psychiatry 54, 1067–1076 (2003).

63. M. Taddei, M. Tettamanti, A. Zanoni, S. Cappa, M. Battaglia, Brain white matter organisation in adolescence is related to childhood cerebral responses to facial expressions and harm avoidance. Neuroimage 61, 1394–1401 (2012).

64. S. D. S. Noordermeer, et al., Structural Brain Abnormalities of Attention-Deficit/Hyperactivity Disorder With Oppositional Defiant Disorder. Biol. Psychiatry 82, 642–650 (2017).

65. H. van Ewijk, et al., The influence of comorbid oppositional defiant disorder on white matter microstructure in attention-deficit/hyperactivity disorder. Eur. Child Adolesc. Psychiatry 25, 701–710 (2016).

66. A. L. Krain, F. X. Castellanos, Brain development and ADHD. Clin. Psychol. Rev. 26, 433–444 (2006).

67. X. Liao, A. V. Vasilakos, Y. He, Small-world human brain networks: Perspectives and challenges. Neurosci. Biobehav. Rev. 77, 286–300 (2017).

68. Z. Cui, et al., Optimization of energy state transition trajectory supports the development of executive function during youth. Elife 9, 1–60 (2020).

