## Supplemental Materials for "Multidimensional brain-age prediction reveals altered brain developmental trajectory in psychiatric disorders"

**Supplementary material**

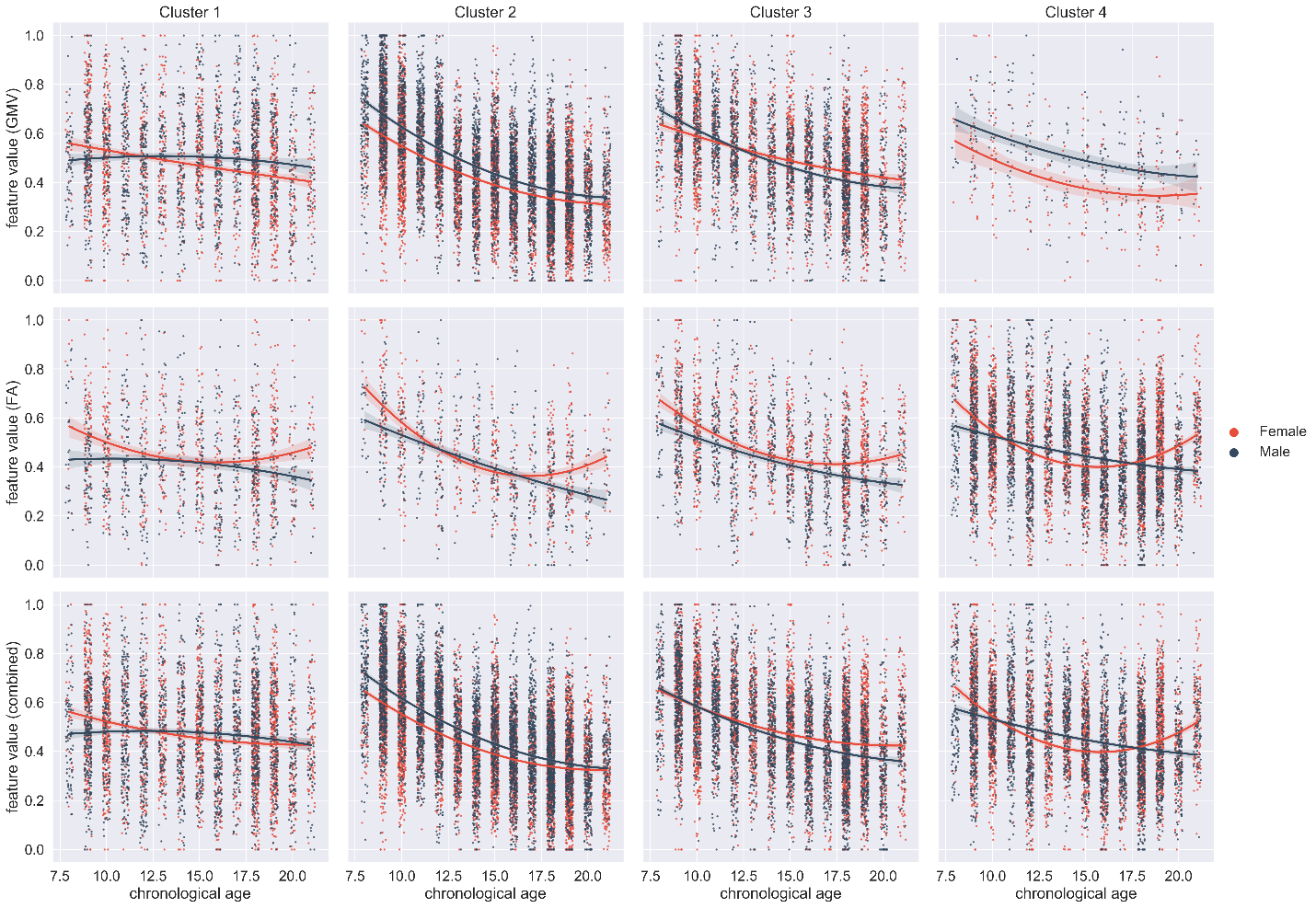

Figure S1. Aggregated developmental trajectories of brain-imaging features in each cluster. Each dot represented a scaled feature value for one brain region of a single subject. Scatterplots in the first and second row showed the aggregated developmental trajectories of the GMV and FA features, respectively. Scatterplots in the third row showed the aggregated developmental trajectories of the GMV and FA features combined. Features in cluster 1 showed a weak linear trend with age and opposite quadratic trends between males and females. Features in cluster 2 showed a strong linear trend and moderate quadratic trend with age. Features in cluster 3 and 4 showed a moderate linear trend. But for cluster 2, the quadratic trend for females was much larger than for males, especially for the FA features. The shaded area represented the confidence interval on the 1,000 bootstrap samples.

Table S1. Brain imaging features in each cluster.

| cluster 1 | cluster 2 | cluster 3 | cluster 4 |
| --- | --- | --- | --- |
| Right Olfactory | Left Precentral | Left Superior Frontal Orbital | Right Superior Frontal |
| Left Parahippocampus | Right Precentral | Right Superior Frontal Orbital | Left Cingulum |
| Right Parahippocampus | Left Superior Frontal | Left Rectus | Right Superior Temporal Pole |
| Right Inferior Occipital | Left Middle Frontal | Right Rectus | Genu.of.corpus.callosum(label) |
| Left Superior Temporal Pole | Right Middle Frontal | Left Anterior Cingulum | Body.of.corpus.callosum(label) |
| Left Middle Temporal Pole | Left Middle Frontal Orbital | Right Anterior Cingulum | Splenium.of.corpus.callosum(label) |
| Right Middle Temporal Pole | Right Middle Frontal Orbital | Left Hippocampus | Inferior.cerebellar.peduncle.L(label) |
| Cerebellum_3_L | Left Inferior Frontal Operculum | Right Hippocampus | Posterior.limb.of.internal.capsule.L(label) |
| Cerebellum_4_5_L | Right Inferior Frontal Operculum | Left Amygdala | Retrolenticular.part.of.internal.capsule.R(label) |
| Cerebellum_4_5_R | Left Inferior Frontal | Right Amygdala | Retrolenticular.part.of.internal.capsule.L(label) |
| Cerebellum_9_R | Right Inferior Frontal | Right Cuneus | Anterior.corona.radiata.R(label) |
| Cerebellum_9_L | Left Inferior Frontal Orbital | Left Superior Occipital | Anterior.corona.radiata.L(label) |
| Cerebellum_9_R.1 | Right Inferior Frontal Orbital | Left Middle Occipital | Superior.corona.radiata.L(label) |
| Cerebellum_10_L | Left Rolandic Operculum | Right Middle Occipital | Posterior.corona.radiata.L(label) |
| Cerebellum_10_R | Right Rolandic Operculum | Left Inferior Occipital | Posterior.thalamic.radiation.R(label) |
| Vermis_1_2 | Left Superior Motor | Left Superior Parietal | Posterior.thalamic.radiation.L(label) |
| Vermis_3 | Right Superior Motor | Right Superior Parietal | Sagittal.stratum.R(label) |
| Vermis_4_5 | Left Olfactory | Left Angular | Sagittal.stratum.L(label) |
| Vermis_9 | Left Superior Medial Frontal | Left Caudate | Cingulum.(hippocampus).R(label) |
| Vermis_10 | Right Superior Medial Frontal | Left Putamen | Cingulum.(hippocampus).L(label) |
| Middle.cerebellar.peduncle(label) | Left Medial Frontal Orbital | Right Putamen | Superior.fronto-occipital.fasciculus.R(label) |
| Fornix.(column.and.body.of.fornix)(label) | Right Medial Frontal Orbital | Cerebellum_Crus1_L | Superior.fronto-occipital.fasciculus.L(label) |
| Medial.lemniscus.R(label) | Left Insula | Cerebellum_Crus1_R | Inferior.fronto-occipital.fasciculus.R(label) |
| Medial.lemniscus.L(label) | Right Insula | Cerebellum_Crus2_L | Inferior.fronto-occipital.fasciculus.L(label) |
| Superior.cerebellar.peduncle.R(label) | Left Middle Cingulum | Cerebellum_Crus2_R | Uncinate.fasciculus.R(label) |
| Superior.cerebellar.peduncle.L(label) | Right Middle Cingulum | Cerebellum_6_L | Uncinate.fasciculus.L(label) |
| Fornix.(cres)./.Stria.terminalis.R(label) | Right Cingulum | Cerebellum_7b_L | Tapetum.R(label) |
| Fornix.(cres)./.Stria.terminalis.L(label) | Left Calcarine | Cerebellum_7b_R | Tapetum.L(label) |
| Cingulum.(hippocampus).R(tract) | Right Calcarine | Cerebellum_8_L | Corticospinal.tract.L(tract) |
|  | Left Cuneus | Vermis_6 | Cingulum.(hippocampus).L(tract) |
|  | Left Lingual | Vermis_8 | Forceps.major(tract) |
|  | Right Lingual | Corticospinal.tract.R(label) | Forceps.minor(tract) |
|  | Right Superior Occipital | Corticospinal.tract.L(label) | Inferior.fronto-occipital.fasciculus.L(tract) |
|  | Left Fusiform | Inferior.cerebellar.peduncle.R(label) | Inferior.fronto-occipital.fasciculus.R(tract) |
|  | Right Fusiform | Cerebral.peduncle.R(label) | Inferior.longitudinal.fasciculus.L(tract) |
|  | Left Postcentral | Cerebral.peduncle.L(label) | Inferior.longitudinal.fasciculus.R(tract) |
|  | Right Postcentral | Anterior.limb.of.internal.capsule.L(label) | Superior.longitudinal.fasciculus.L(tract) |
|  | Left Inferior Parietal | Posterior.limb.of.internal.capsule.R(label) | Uncinate.fasciculus.L(tract) |
|  | Right Inferior Parietal | Posterior.corona.radiata.R(label) | Uncinate.fasciculus.R(tract) |
|  | Left Supramarginal | External.capsule.R(label) | Superior.longitudinal.fasciculus.L(tract) |
|  | Right Supramarginal | External.capsule.L(label) | Superior.longitudinal.fasciculus.R(tract) |
|  | Right Angular | Superior.longitudinal.fasciculus.R(label) |  |
|  | Left Precuneus | Anterior.thalamic.radiation.L(tract) |  |
|  | Right Precuneus | Anterior.thalamic.radiation.R(tract) |  |
|  | Left Central Paracentral Lobule | Corticospinal.tract.R(tract) |  |
|  | Right Central Paracentral Lobule | Superior.longitudinal.fasciculus.R(tract) |  |
|  | Right Caudate |  |  |
|  | Left Pallidum |  |  |
|  | Right Pallidum |  |  |
|  | Left Thalamus |  |  |
|  | Right Thalamus |  |  |
|  | Left Heschl |  |  |
|  | Right Heschl |  |  |
|  | Left Superior Temporal |  |  |
|  | Right Superior Temporal |  |  |
|  | Left Middle Temporal |  |  |
|  | Right Middle Temporal |  |  |
|  | Left Inferior Temporal |  |  |
|  | Right Inferior Temporal |  |  |
|  | Cerebellum_6_R |  |  |
|  | Vermis_7 |  |  |
|  | Anterior.limb.of.internal.capsule.R(label) |  |  |
|  | Superior.corona.radiata.R(label) |  |  |
|  | Cingulum.(cingulate.gyrus).R(label) |  |  |
|  | Cingulum.(cingulate.gyrus).L(label) |  |  |
|  | Superior.longitudinal.fasciculus.L(label) |  |  |
|  | Cingulum.(cingulate.gyrus).L(tract) |  |  |
|  | Cingulum.(cingulate.gyrus).R(tract) |  |  |
